# In vitro inhibition of the CFTR ion channel in the *Macaca mulatta* cervix thickens cervical mucus

**DOI:** 10.1101/2024.10.14.618249

**Authors:** Rachel J. Looney, Mackenzie Roberts, Matthew Markovetz, Rachelle Godiah, Shan Yao, Kirsti Golgotiu, Shuhao Wei, Chris Cellucci, Leo Han

**Affiliations:** Division of Reproductive and Developmental Sciences, Oregon National Primate Research Center, Beaverton, OR, USA; Marsico Lung Institute, University of North Carolina, Chapel Hill, NC, USA; Department of Biomedical Engineering, Oregon Health and Science University, Portland, OR, USA; Department of Obstetrics and Gynecology, Oregon Health and Science University, Portland, OR, USA

**Keywords:** CFTR, Cervix, Mucus, Fertility, Inh-172, ASL, PTMR

## Abstract

Cervical mucus changes throughout the menstrual cycle in response to hormonal fluctuations, regulating access of sperm and pathogens to the reproductive tract. CFTR is an anion channel that plays a critical role in mediating epithelial mucus secretions. Primary endocervical cells obtained from rhesus macaques *Macaca mulatta* were cultured using conditional reprogramming and treated with vehicle controls or CFTR inhibitors. In order to measure changes in hydration and viscosity of secreted mucus, we adapted two airway mucus assays, airway surface liquid and particle-tracking microrheology, for our endocervical culture system. Endocervical cells treated with CFTR inhibitors demonstrated dehydrated, thicker mucus secretions compared to controls in both assay outputs. Our studies suggest that CFTR may be an important mediator of fertility changes and provide experimental evidence for the infertility phenotype seen in women with cystic fibrosis. Additionally, assays developed in these studies provide new endpoints for assessing cervical mucus changes in vitro.

**Summary Sentence:** Inhibition of CFTR, a key epithelial ion channel in primary endocervical cells leads to dehydrated, thickened mucus production in vitro

## Introduction

Epithelial cells, lining the endocervical canal, control sperm and pathogen entry into the female reproductive tract through changes in cervical mucus secretion. Over the course of the menstrual cycle, fluctuations in ovarian sex-steroids, estradiol (E2) and progesterone (P4), regulate the biophysical properties of secreted mucus[1,2]. When E2 levels peak during the ovulation window, cervical mucus increases in quantity and hydration, and decreases in viscosity, creating a permeable gel that allows sperm to move through the cervix and into the upper tract. Under the influence of P4, following ovulation, cervical mucus increases in concentration, becomes more viscous, and subsequently blocks sperm from entering the reproductive tract [3].

The mucus produced by the cervix is a heterogenous mixture of water, mucin and non-mucin proteins, lipids, carbohydrates, and ions. Outside the cervix, other epithelia, including the lung, GI tract, and eye, also produce mucus that maintain organ health and functions. Mucins and proteins found in mucus are retained across organ systems [4]. The Cystic Fibrosis Transmembrane Conductance Regulator (CFTR) ion channel has been shown to be a critical regulator of mucus characteristics in all mucus secreting epithelia. Mutations in the CFTR gene lead to dysfunctional channel activity and the autosomal recessive disease, cystic fibrosis (CF). CF is characterized by systemically thickened mucus secretions that can cause multi-organ dysfunction in the airway, GI tract, and reproductive tract. In the female reproductive tract, CF is notable for thickened endocervical mucus as well as sub-fertile and infertile phenotypes [5,6].

The CFTR channel moves chloride and bicarbonate (ions) into the lumen, promoting hydration and the thinning of mucus through two primary mechanisms (Figure 1). First, CFTR maintains an ionic gradient for water to move osmotically [7]. A second channel, the epithelial sodium channel (ENaC), moves sodium into the cell, from the lumen. Thus, increased CFTR activity coupled with decreased ENaC, results in net sodium and chloride ion retention in the lumen and increased water movement to the lumen, with the end result of hydrating mucus and increasing viscosity. Conversely, decreased CFTR activity and/or increased ENaC activity will have the opposite effect of removing salt from the lumen and dehydrating mucus. The second mechanism specifically relates to the critical role CFTR plays in mucin protein unfolding. Large mucin glycoproteins are tightly packaged intracellularly before their release. This is accomplished by storing mucins in secretory granules with low pH and high amounts of cations (e.g., calcium) to shield the negatively charged carbohydrate-rich ends from themselves while they are stored. Upon release, the higher luminal pH from CFTR secreted bicarbonate, as well as the presence of other luminal anions remove these cations resulting in the immediate expansion and unfolding of the mucins by as much as 500 to 1000-fold in size. In their expanded form, the now “opened” mucin branches are fully exposed and ready to bind water [8]. Thus, when CFTR is dysfunctional, mucins are also unable to fully expand and bind water, leading to the high density, low hydration, and high viscosity mucus seen in CF [9].

**Figure 1:**
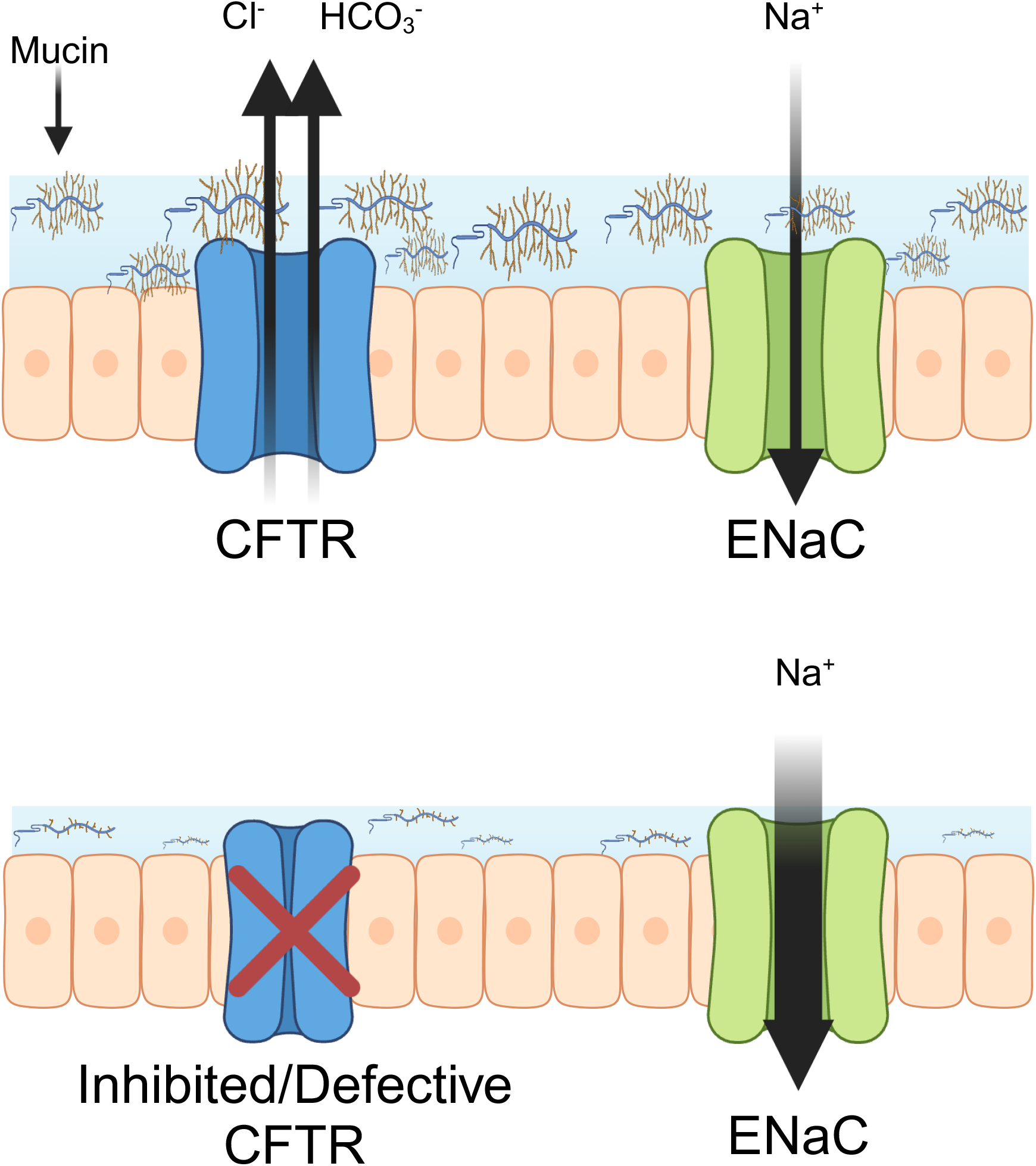
This figure illustrates the relationship between CFTR and ENAc as it pertains to regulating mucus concentration. **Alt text:** A figure where two ion channels are seen to regulate mucus concentration. The flow of ions can also be seen.

Our prior studies demonstrated that CFTR in the cervix is both functionally and hormonally regulated [10]. While these findings may suggest that CFTR plays a role in regulating mucus changes in the cervix, CFTR inhibition of endocervical cells and its effect on mucosal thickness and hydration has yet to be demonstrated in an experimental model. Further elucidating the contribution of CFTR in cervical mucus regulation would more precisely define its potential as a target for both fertility and contraceptive studies in the cervix. However, in vitro models for studying endocervical mucus secretion lack experimental endpoints for measuring mucus changes in response to perturbation experiments. To overcome this limitation, we adapted from airway studies of mucus, airway surface liquid (ASL), as a method for measuring mucus hydration in vitro. In vitro ASL correlates tightly to interfacial hydration in vivo [11–13](Figure 2A). ASL in normal airway epithelia [11,14] is lost in CF cells due to mucosal dehydration [15] and can be restored in mutant cell lines with novel CFTR modulator therapies [13,16]. While ASL can be directly measured using confocal microscopy in the XZ axis [11], ASL can also be assessed by measuring the edge-concave meniscus formed under brightfield microscopy that allows 96-well experimental measurement and replication [12](Figure 2B). This measurement of ASL and can also then be used for high-throughput studies of CFTR modulator therapies [16].

**Figure 2:**
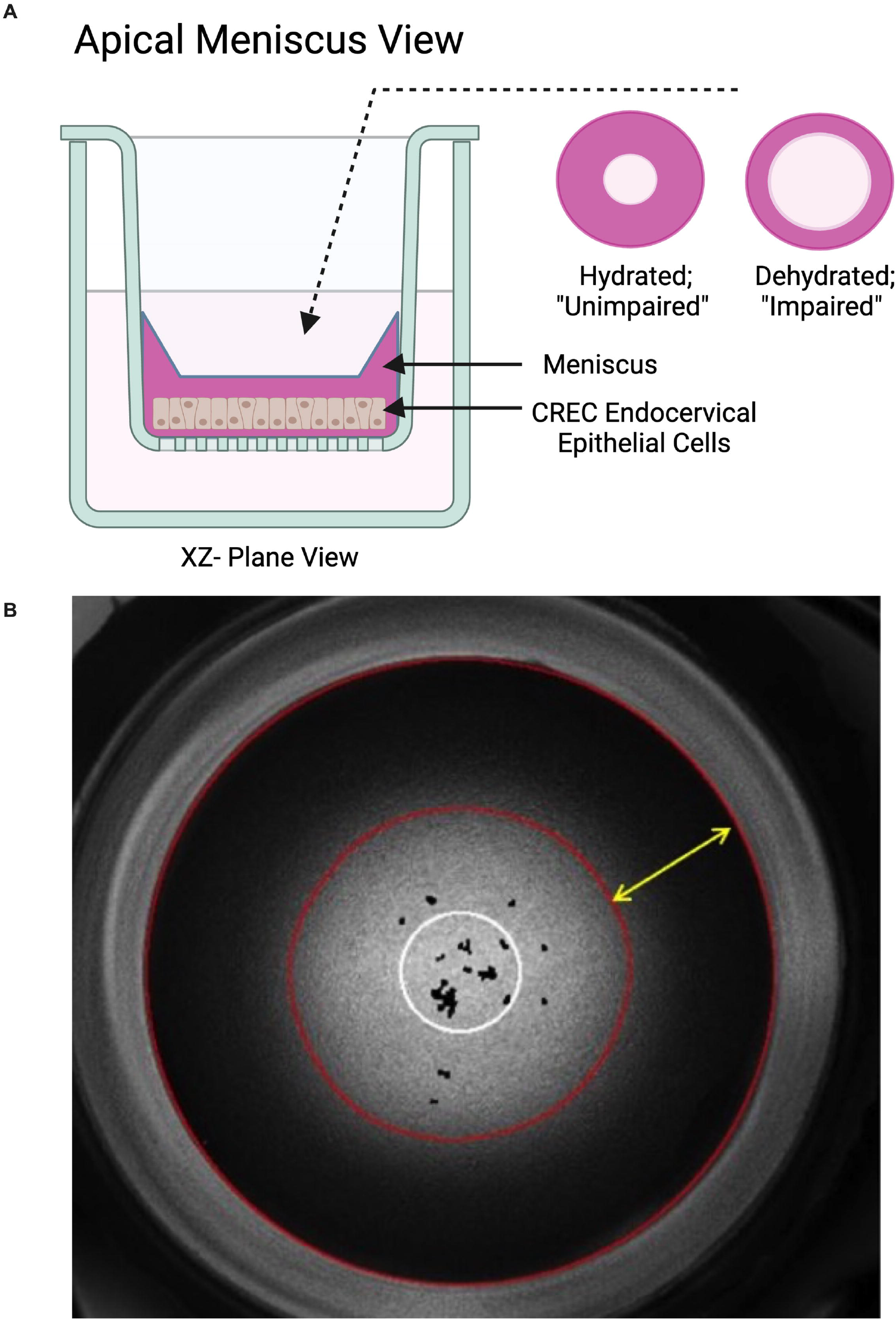
(A) This figure illustrates the XZ-plane of a Transwell® during an experiment. The apical meniscus view shows the change in meniscus corresponding to CFTR function. (B) This figure demonstrates the image obtained from brightfield microscopy used in ASL studies. The cells can be seen in the center of the field forming a monolayer. The darker outer ring is the meniscus or annulus formed by mucus and PBS on the apical surface of the Transwell®. **Alt text:** (A) A cross-sectional diagram showing an experimental setup of endocervical epithelial cells) within a liquid-filled container. To the right of the container, there are two circular illustrations labeled “Apical Meniscus View.” One circle represents a hydrated, “unimpaired” meniscus, while the other shows a dehydrated, “impaired” meniscus. (B) A grayscale circular image resembling a microscope view, featuring concentric rings with distinct boundaries outlined in white. The distance between edge of the well and the lighter region is the annulus.

While ASL is a measure of liquid-ion transport functionality, it does not provide a direct readout of the biophysical status of the mucus in the lumen. Multiple studies have shown that mucus concentration is the primary driver of aberrant biophysical properties (i.e. viscoelasticity) observed in CF [17,18]. This has been demonstrated with classical rheometric and microscale techniques relevant to the biophysics experienced by sperm within mucus, generally termed particle tracking microrheology (PTMR). PTMR exploits the thermal diffusion of small (~1 μm) fluorescent probes to sense the viscoelasticity of the mucus network. Thus, when volumes of a liquid or gel, are too small for normal rheological instruments, they can be assessed with PTMR.

In this investigation, we adapted both ASL measurements of mucus hydration and PTMR measurements of viscosity to our endocervical cell culture system and used them to assess changes in mucus secretion in response to hormonal treatments and CFTR inhibition. We hypothesized that selectively inhibiting CFTR with several known small molecule inhibitors would result in decreased mucosal hydration, and increased mucus viscosity in vitro.

## Methods

### Primary cell culture

Rhesus macaque cervical tissue was obtained from the Oregon National Primate Research Center tissue distribution program. Tissue was obtained from animals ranging from 4-15 years with and without history of prior vaginal deliveries. Endocervical epithelia were scalpel separated from the cervical stroma, enzymatically digested, and cultured using conditionally reprogramming conditions as previously described [19]. Expanded cells were passaged at 80% and placed onto HTS Transwell® 96-well and 12-well Permeable Polyester (PET) Supports (0. 4 μm)(Corning, New York) in a serum-free media (ReproLife CX Lifeline Cell Technology, Frederick, Maryland), supplemented with 1M calcium chloride and 10nM E2 and taken to an air-liquid interface (ALI).

### Airway surface Liquid

We used transepithelial electrical resistance (TEERs) to determine when cells were ready for experimentation, as TEER levels assess tight junction integrity, cellular polarization and differentiation[20]. After 10-14 days at an air-liquid interface (ALI), when TEER>300 Ω*cm^2^, we performed the airway surface liquid assay (ASL) on treatment naïve cells[19]. For ASL, we adapted an assay variation used for high-throughput screening (HTS) in human bronchial epithelial (HBE) cultures that measures the cellularity ability to resorb excess saline added to the apical (mucus) side of ALI culture [16]. Both the rate and magnitude of change to the ASL layer, as the cells try to return the overlying ASL to homeostasis, reflect the underlying mucus secretion and resorption properties of the epithelia. We developed two different versions of the resorption assay. In a version similar to the previously published HTS, phosphate-buffered saline (PBS; 8 *µ*L per well) (Thermo-Fisher Scientific) was added to the apical surface, forming a concave meniscus, and images were taken using brightfield microscopy (2x PlanApoλ NA 0.10 magnification, BZ-X Series, Keyence, Itasca, Illinois). We additionally assessed these changes using an apical addition of 8 *µ*L per well of 100 mM mannitol in PBS. Mannitol is a sugar alcohol that demonstrates osmotic based expansion of ASL in HBE cells and has been studied in clinical trials as potential treatment for CF lung disease [21,22]. We used this combined method of assessing ASL (mannitol+ PBS) as we found that it more robustly defined changes between controls and treatment in ASL studies [23]. Images were taken pre-treatment, immediately post-treatment (0 hours), 24, 48, 72, and 96 hours.

Following image acquisition, we used custom FIJI [24] scripts to automatically calculate the change in fluid meniscus. The scripts use pixel thresholding to determine the size of the meniscus by measuring the annulus; the annulus is the area of dark fluid meniscus that emanates from the edge of the well to the lighter center region (Figure 2B). This area increases with greater apical hydration and decreases with reduced hydration. The metrics were reported as magnitude change from baseline (time=0 hours) in downstream analysis and statistics. More negative values correspond to greater reduction in the meniscus from the 0Hr time point (time of apical application of PBS). Thus, larger change in meniscal reduction indicates increased dehydration of the cell. As part of our custom scripts, we additionally adjusted for overlying artifacts (i.e. overlaying cellular mucus or debris) in the cell cultures using thresholding, and automatic identification of outliers based on a pre-determined 95% confidence interval. Using the 96-well plate, treatment conditions were repeated using two different plates to account for edge effects with a total of either 48 or 96 wells per condition. Experiments represent at least three or more unique primary cell lines. We used cells between passage two and five. *Particle Tracking Microrheology*

To use PTMR for measuring viscosity, 1 µm diameter carboxylated FluoSpheres™ (ThermoFisher, Fremont, CA), henceforth referred to as “beads”, were added to the lumen of cell cultures. The thermally driven motion of the beads was tracked to determine the viscoelastic properties of the luminal mucus. Bead motion was recorded for 10s at a rate of 60 frames/s using a 40x air objective on a Zeiss AxioVert A1 Fluorescent microscope (Zeiss, Jena, Germany). Individual bead trajectories were measured automatically using a custom Python program that uses TrackPy [25] for bead localization and tracking. Bead motion was converted into mean squared displacements (MSD) and complex viscosity (η*) values in accordance with the mathematics described previously [25,26].

### CFTR Inhibition

We used two different small molecule CFTR inhibitors: Inh-172 (5 or 10*µ*M) or GlyH-101 (5*µ*M) (MilliporeSigma, Burlington, Vermont)[27]. Inh-172, a thiazolidinone-class CFTR inhibitor, binds inside the channel pore near a transmembrane helix, stabilizing the closed channel conformation, and producing a CFTR mutant like phenotype in inhibited cells[28,29]. GlyH-101 is a glycine hydrazide-class inhibitor that also binds to the membrane-spanning pore region, blocking the channel. Both inhibitors were chosen because they are widely used for CFTR inhibition experiments, do not affect cell viability at effective concentrations, and their mechanism of action and channel specificity are well documented[28–31]. Treatment was applied to the basolateral compartment of the Transwell® every other day.

### Statistics

One-way ANOVA was administered using SAS statistical software (Cary, NC). A general linear model testing least squares means for effect was generated and p-values less than 0.05 were considered significant. “N” was denoted as the number of primary cell cultures used for 96-well experiments. Magnitude or reduction in the meniscus from baseline are expressed at means ± SEM.

## Results

### Airway Surface Liquid Baseline Assays

We began by demonstrating the ability of our assay to measure changes in apical hydration using mannitol only treatments [32]. Cells were treated with vehicle or mannitol (100mM and 300mM) in PBS (8 *µ*L) applied apically on day 1. Mannitol treated cultures demonstrated persistently elevated ASL meniscus measurements at all time points (p<.05, Figure 3). This effect was dose dependent with increased ASL at 300 mM compared to 100 mM (Figure 3).

**Figure 3:**
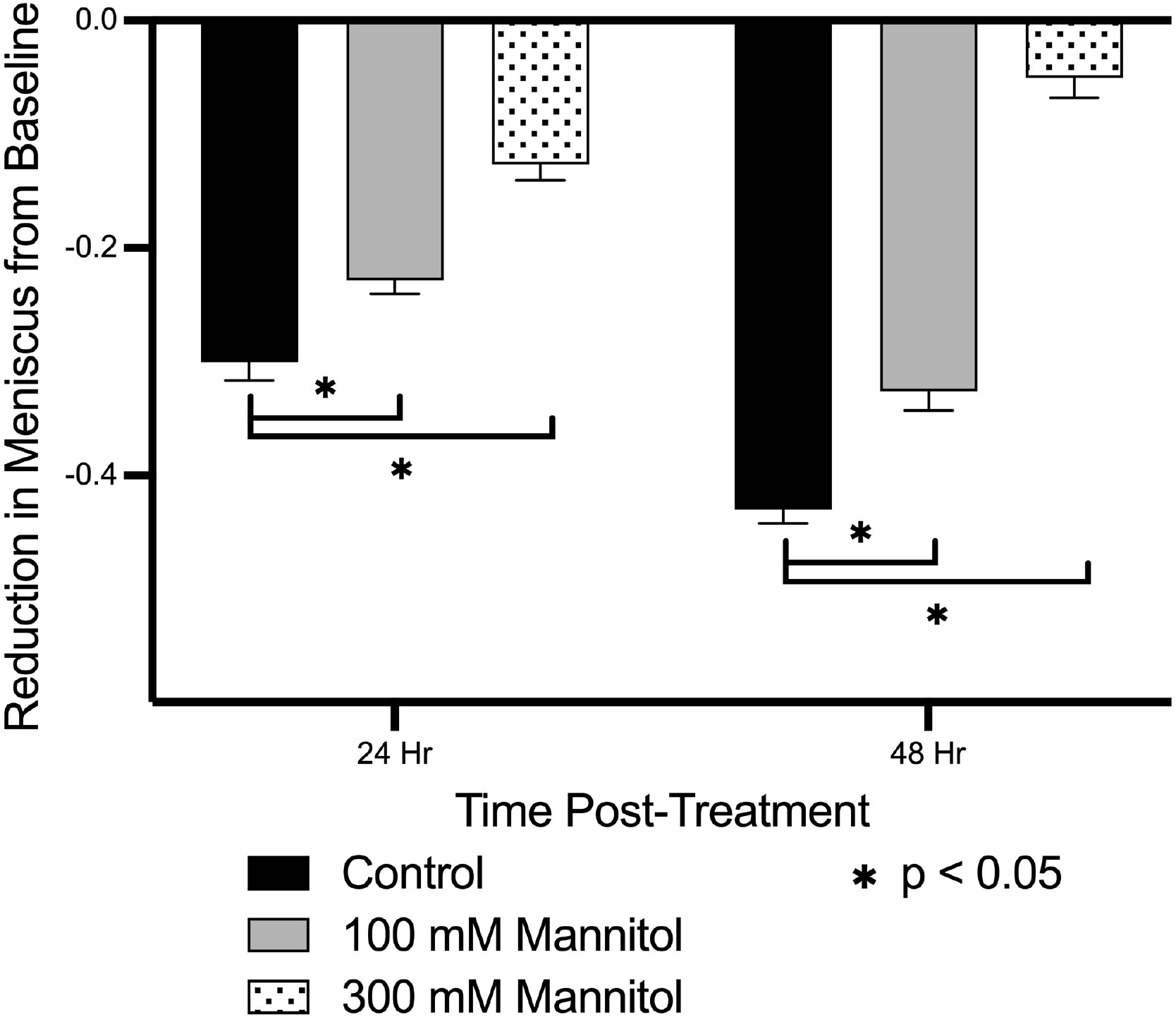
Effect of mannitol on ASL hydration. Across all time points (24, 48 hours from administration) mannitol (100 mM and 300 mM) reduced the measured change in meniscus from baseline compared to the control (n=4, P<0.05). **Alt text:** Bar chart showing reduction in meniscus from baseline for various treatment groups over time.

We also sought to demonstrate that ASL measurements responded to hormonal treatment of mucus. Our prior studies demonstrate that conditionally reprogrammed endocervical cells maintain estrogen receptor (ER) and progesterone receptor (PGR) expression and hormonal sensitivity[19,33].

Clinically, E2 peaks during ovulation and results in thinner, hydrated mucus. During the luteal phase, P4 dominates, and mucus becomes thicker and more dehydrated [34,35]. Thus, we would expect E2 to increase ASL relative to control, and P4 to decrease ASL relative to control [3]. To do this, we treated differentiated cells with E2 (10 nM) or E2/P4 (10 – 14 days of 10 nm E2 priming followed by 1nM E2 and 100 nM P4) basolaterally and compared the responses to the vehicle only controls [36–38]. Treatment with 10 nM E2 resulted in greater ASL hydration when compared to the control condition (p<0.001) (Figure 4A). Similarly, treatment with 100nM P4 resulted in greater reduction in ASL than 10 nM E2 at every time point (P<0.05) (Figure 4B).

**Figure 4:**
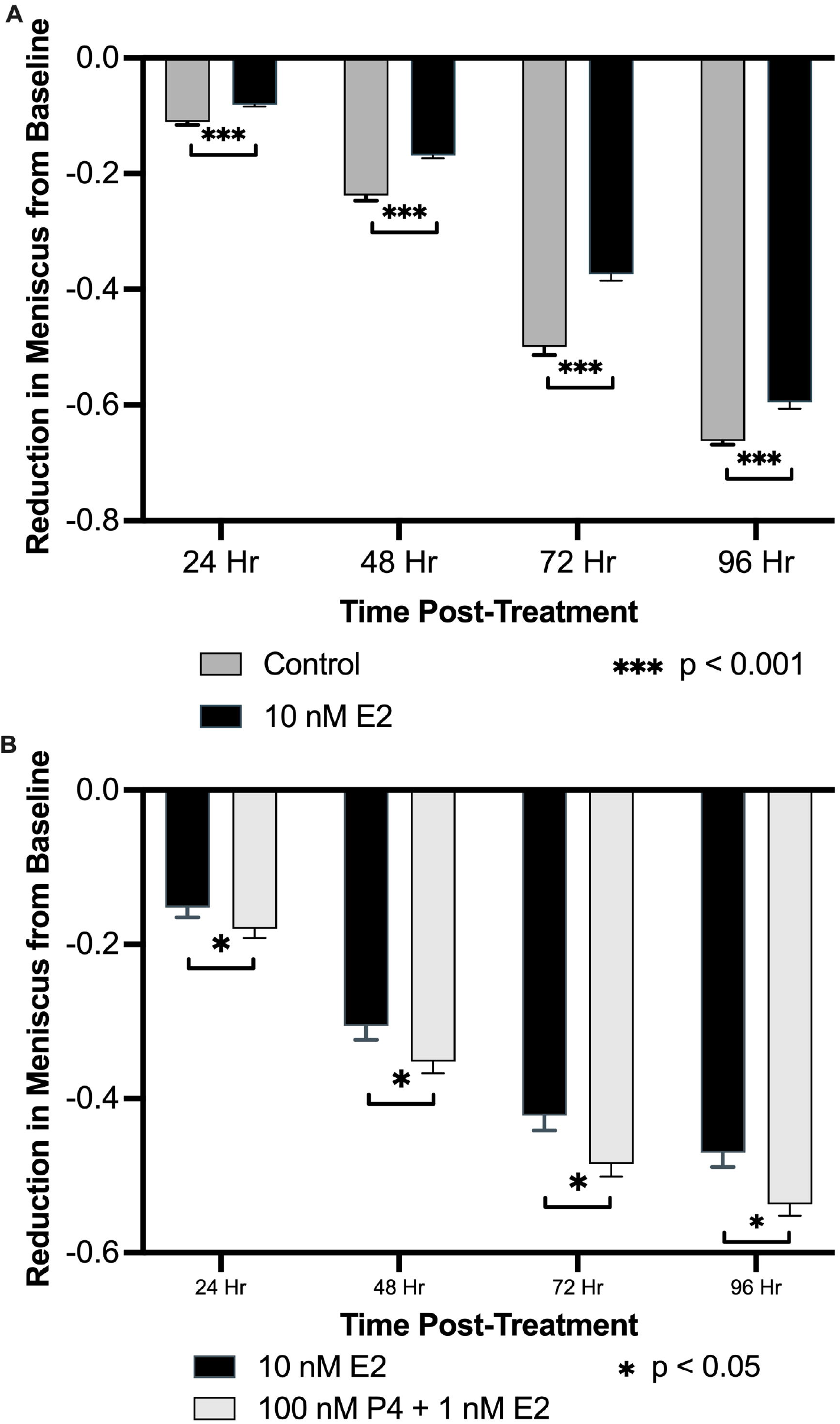
(A). Effect of E2 on ASL hydration. Across all time points (24, 48, 72, 96 hours from administration) E2 (10 nM) reduced the measured change in meniscus from baseline compared to the control(n=6, P<0.001). (B). Effect of E2 and P4/E2 on ASL hydration. Across all time points (24, 48, 72, 96 hours from administration) 100 nM P4 (after E2 priming) increased the measured change in meniscus from baseline as compared E2 only (n=6, P<0.05). **Alt text:** Bar charts showing reduction in meniscus from baseline for various treatment groups over time.

### ASL response to CFTR Inhibition

10*µ*M Inh-172 resulted in statistically significant ASL dehydration by 48, 72, and 96 hours (p<0.01), while 5*µ*M GlyH-101 had a smaller but still significant dehydration effect (p<0.05), compared to the control condition (Figure 5A, 5B). Using mannitol in conjunction with PBS amplified the dehydration response across all time points by up to three-fold compared to PBS alone. (Figure 5B).

**Figure 5:**
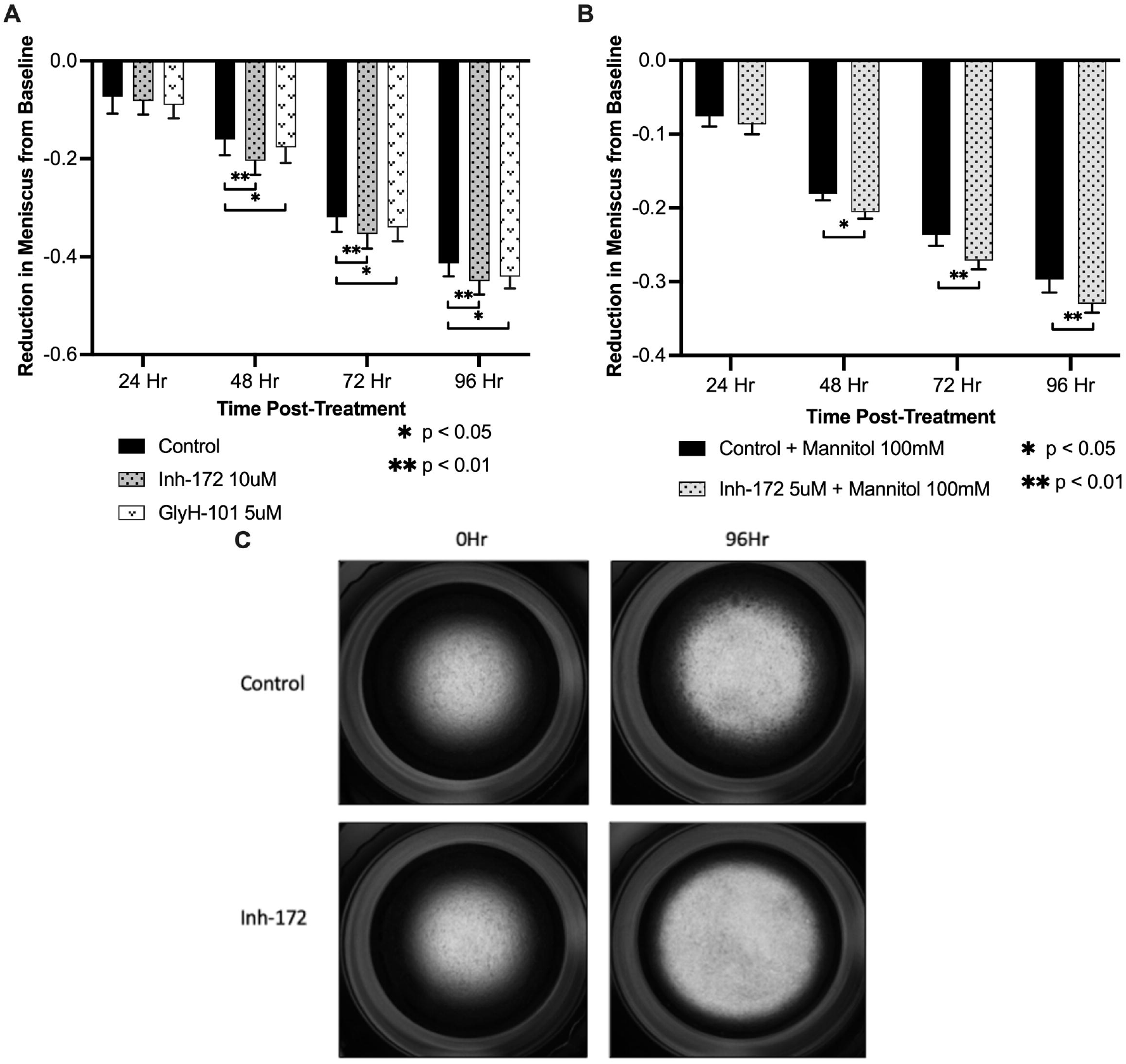
(A) Effect of InH-172 and GlyH-101 on ASL Hydration. Across time points 48, 72, and 96 hours from administration, 10 μM Inh-172 (n=6) and 5 μM GlyH-101(n=6) increased the measured change in meniscus from baseline compared to the control (P<0.01, P<0.05 respectively). (B) Effect of Inh-172 and Mannitol on ASL Hydration. Across time points 48, 72 and 96 hours from administration, 5 μM Inh-172 with 100 mM mannitol (n=6) increased the measured change in meniscus from baseline compared to the control (P<0.05, P<0.01, P<0.01 respectively). (C) Representative images obtained at 0 hr and 96 hr with control and Inh-172. These examples were chosen as visually greater reduction of the meniscus in the Inh-172 condition is seen. **Alt text:** Bar charts showing reduction in meniscus from baseline for various treatment groups over time. Chart A compares three groups, and chart B compares two groups. A collection of four images depicting a single culture well. The dark region represents the meniscus which decreases over time in both conditions.

### PTMR response to CFTR-inhibition

The reduction of ASL volume by Inh-172 was visually notable (Figure 5C), resulting in increased mucus concentration and, therefore, an increase in mucus viscoelasticity. This was quantified by PTMR, which demonstrated decreased bead diffusion three-fold, resulting in visually smaller diffusive trajectories (Figure 6). Accordingly, mucus viscoelasticity increased significantly after Inh-172 treatment from 0.15 Pa·s to 0.48 Pa·s in n=6 cultures (p<0.01)(Figure 7) and had a more uniform viscosity than untreated controls.

**Figure 6:**
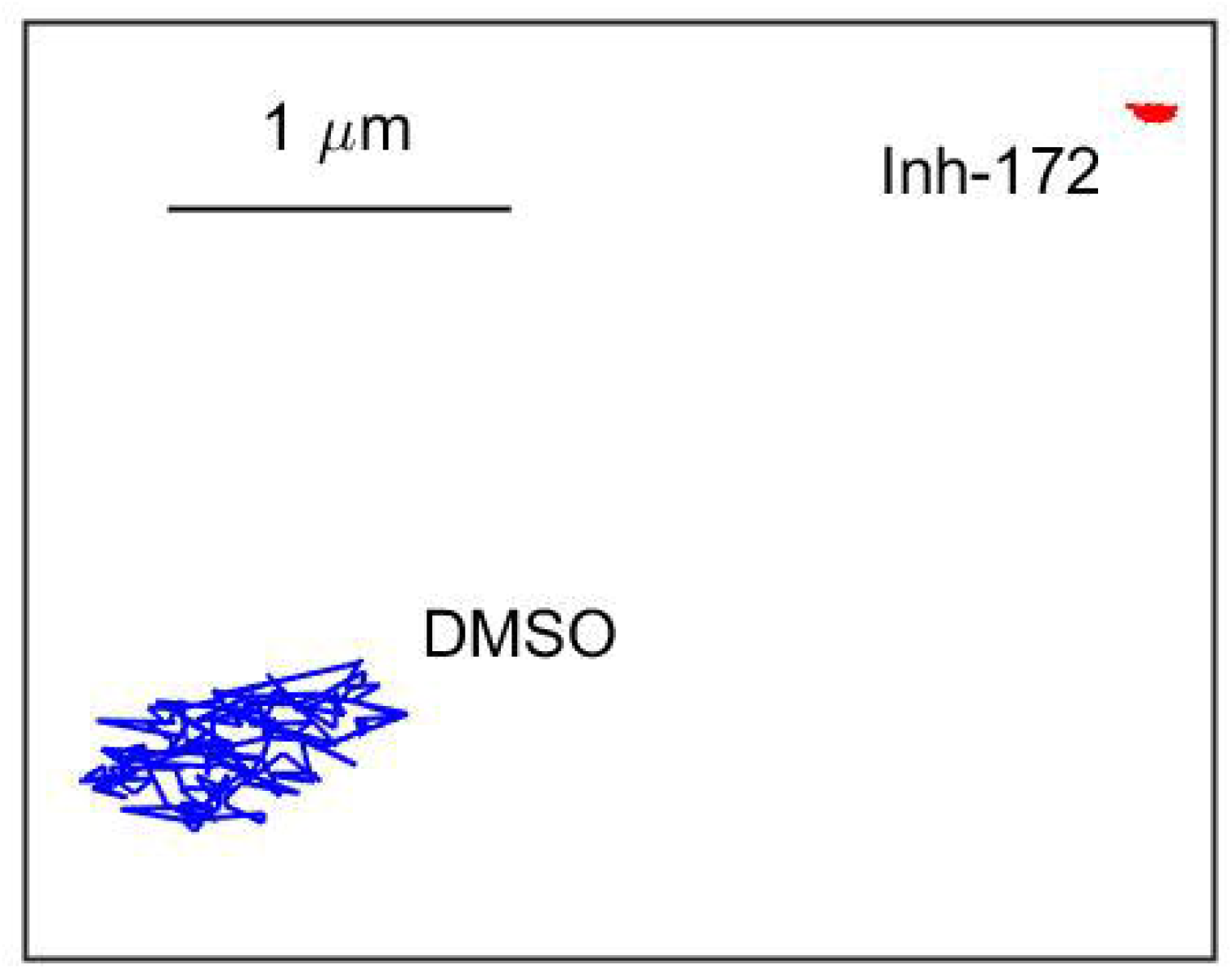
Representative bead trajectories after treatment with either DMSO or Inh-172. Treatment with Inh-172 resulted in significantly less bead movement compared to DMSO treatment. **Alt text:** a figure representing the movement of microscopic beads in mucus secreted by cells treated with DMSO (blue) and Inh-172 (red).

**Figure 7:**
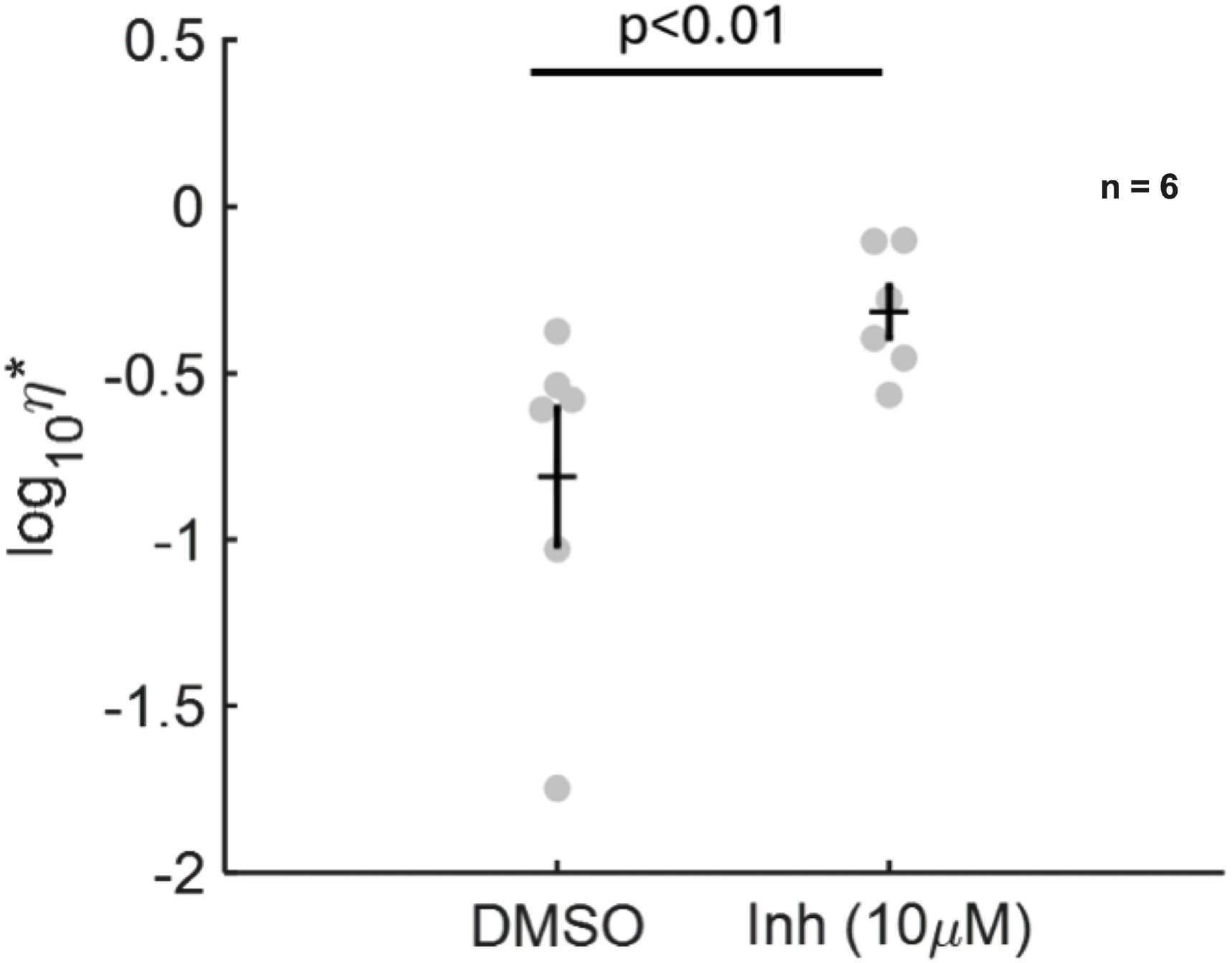
Effect of 10 μM Inh-172 on mucus complex viscosity (η*). Treatment with Inh-172 for 72 hours increased mucus secretion η* (p<0.01; n=6). **Alt text:** a graph demonstrating the difference between DMSO and Inh-172 on complex viscosity.

## Discussion

Using two different phenotypic assays measuring mucus properties, we demonstrate that inhibition of CFTR in vitro changes secreted mucus by endocervical epithelial cells. Specifically, our results are consistent with the *clinical* findings that mucus is more hydrated in response to E2, dehydrated when exposed to P4, and that CFTR inhibition results in abnormal, dehydrated, thickened mucus secretions similar to that seen in cystic fibrosis [6]. Combined with prior studies that demonstrate the hormonal regulation of CFTR during the menstrual cycle, these findings further support the integral role of CFTR in endocervical mucus changes.

Additionally, this study translates two assays, airway surface liquid (ASL) and particle tracking microrheology (PTMR), into endocervical mucus studies [12,25]. New to endocervical mucus studies, these assays provide experimental endpoints that directly reflect mucus changes secreted by cultures, correlated to biophysical properties of mucus itself. ASL, in particular, has potentially high experimental utility as it has been both well validated in mechanistic studies of mucus regulation and used in a high-throughput fashion for drug screening [16]. PTMR is a useful technique for this study for two reasons: it operates and measures forces at the same scale as sperm in the cervical mucus environment, and it has been shown that there is a strong correlation between in vitro mucus models and in vivo mucus in the airway [25]. Uniquely, the reduced ciliary activity/presence in the conditionally reprogrammed endocervical cell cultures allows for PTMR to be performed *in situ*, further increasing its utility here.

Both assays provide potential endpoints for future studies of mucus in vitro. One challenging characteristic of studying endocervical mucus is the lack of valid small animal models. Classic mammalian models, such as rodents, are poor options for studying cervical mucus changes or lower tract fertility as they lack a mucus secreting endocervix and sperm is ejaculated directly into the uterus [39]. In larger mammalian models, experimental rigor is challenging due to biological variability, semi-quantitative nature of mucus appraisal, and invasiveness for mucus collection. An in vitro model provides a tool for mechanistic studies to determine the key regulators of mucus changes such as CFTR, but also a platform for drug discovery. With recent advances in primary cell culture such as conditional reprogramming, there is further potential for study of genetic differences and personalization of drugs response [40].

Our previous studies demonstrate that CFTR is a hormonally regulated ion channel with E2 (mid cycle) conditions upregulating CFTR gene expression and, conversely, P4 (luteal) conditions downregulating CFTR expression [10]. In CF, the CFTR gene mutation produces a defective channel, resulting in thick, viscous cervical mucus. Prior to the drug discovery and development of small molecules that modulate mutant CFTR function, women with CF had known infertility/subfertility phenotypes that were largely attributed to these mucus changes, preventing sperm penetration [41]. In our study, we used inhibitors to mimic CFTR mutation, recapitulating this phenotype in vitro. Thus, our study is one of the first to experimentally confirm the relationship of CFTR inhibition and mucus changes using loss of function experiments in vitro. Additionally, it provides evidence that CFTR is likely a critical mediator of mucus changes in normal menstrual cycles.

Our in vitro study does not fully recapitulate either tissue level characteristics of epithelial biology nor the full content of mucus secretions produced in vivo. However, the conditionally reprogrammed platform used in this investigation provides a robust model where we can investigate epithelial changes under multiple treatment conditions, while evaluating mucus production and quality. We have previously demonstrated in proteomic studies, high similarity between the mucus produced in vitro using this cellular system to in vivo mucus [10,19,42], and epithelial only cell models have been used to high success in disease investigations and drug discovery in other mucosal systems [40,43].

The extent to which CFTR is responsible for mucus changes in the cervix is not fully quantified by our studies. CFTR, while a critical channel with a well-known mutated human phenotype, is part of a large ecosystem of ion channels. There are other ion channels, such as the epithelial sodium channel (ENaC) that are hormonally active and may be involved in mucus hydration [44]. Investigating the relative contributions of different ion channels in mucosal hydration using our endpoints could provide a more comprehensive understanding of female fertility and the development of nonhormonal drug therapies.

Additionally, future studies might develop our findings further by examining the effect of CFTR inhibition on fertility and mucus consistency and hydration in vivo using a whole animal model.

## Conclusion

Cervical mucus produced by the epithelia in the endocervix is a hormonally induced regulator of female fertility. In this investigation we validate and employ two different techniques, ASL and PTMR, to measure the viscosity and hydration of endocervical cell mucus. We determined that the inhibition of CFTR using small molecule inhibitors results in significant dehydration and increased viscosity of cervical mucus in vitro. These findings are consistent with the physiology seen clinically and predicted by molecular studies of CFTR gene expression.

## Author contribution statement

RL: performed experiments, analyzed data, wrote paper

MR: performed experiments, analyzed data, wrote paper

MM: performed experiments, analyzed data, wrote paper

RG: performed experiments, analyzed data, wrote paper

SY: performed experiments, analyzed data

KG: performed experiments, analyzed data, wrote paper

SW: performed experiments, analyzed data

CC: performed experiments, analyzed data

LH: conceived the study, performed experiments, analyzed data and wrote the paper

## Acknowledgements

This work was supported, in whole or in part, by the Bill & Melinda Gates Foundation [INV-024195]. Under the grant conditions of the Foundation, a Creative Commons Attribution 4.0 Generic License has already been assigned to the Author Accepted Manuscript version that might arise from this submission.

## Data Availability Statement

The data underlying this article will be shared on reasonable request to the corresponding author.

